# Enhanced Fidelity and Sequence Generality of an RNA Polymerase Ribozyme

**DOI:** 10.64898/2026.10.18.759947

**Authors:** David P. Horning, Samantha Nasser, Gerald F. Joyce

## Abstract

Directed evolution has been used to develop RNA enzymes (ribozymes) that catalyze the RNA-templated polymerization of RNA. The more advanced polymerases can synthesize long RNA products, but have shortcomings with regard to copying accuracy and template sequence generality, which are impediments to using the polymerase to replicate and evolve other functional RNAs. By applying stringent selection pressure over the course of 33 successive rounds of evolution, both the fidelity and sequence generality of the polymerase were greatly improved. When synthesizing large functional RNAs such as the class I ligase ribozyme, the fidelity improved from 86% to 94% per nucleotide, due to both reduced incorporation of mismatched nucleotides and reduced propensity to extend partial-length products that have mismatched termini. The improved polymerase is also able to synthesize a broad range of RNA products, including those that require copying templates that are highly structured or contain sequences that previously were difficult to copy.

## Introduction

RNA has been proposed as the first informational macromolecule of life, prior to the emergence of genetically encoded proteins, with RNA fulfilling the roles of gene and enzyme played by DNA and protein in modern life.^1–4^ In this “RNA world”, the primary means by which RNA genes would have been propagated is through RNA-catalyzed, RNA-templated synthesis of RNA. Some functional RNA elements from this era, most notably the ribosome, still exist in extant life. However, there is no known biological RNA that catalyzes RNA-templated polymerization of RNA. Instead, directed *in vitro* evolution has been used to develop RNA polymerase ribozymes, starting from libraries of random-sequence RNAs.^5^ The most advanced of these polymerases can copy other RNAs, including other ribozymes.^6,7^

The level of genomic complexity that was reached in the RNA world would have depended on both the sequence generality and accuracy of the genome copying machinery. Sequence generality refers not just to the primary sequence of the RNA being copied, but also to its structure. A well-defined secondary and tertiary structure are essential for RNA’s role as a functional molecule, but are an impediment to a polymerase that must operate on an unfolded, linear template. The fidelity of copying determines the amount of genomic information that can be propagated over the generations. If the error rate exceeds a critical threshold, selection cannot maintain genetic information against the mutational load, and the population decays toward random sequences.^8^ Even for copying a small ribozyme of ∼50 nucleotides, the theoretical threshold corresponds to approximately 97% copying fidelity per nucleotide, although actual thresholds can be substantially more relaxed because neutral and compensatory mutations can preserve function.^9^

The most active and general polymerase ribozymes discovered to date derive from the class I ligase ribozyme, first isolated from a population of random-sequence RNAs containing 220 nucleotides.^5,10^ Early directed evolution efforts sought to convert the ligase to a polymerase that can extend an RNA primer on a complementary RNA template by incorporating all four nucleoside 5’-triphosphates (NTPs). These efforts resulted in an RNA-dependent RNA polymerase that can add up to 14 nucleotides,^11^ and further evolution increased the length of extension to many dozens of nucleotides.^6^ The resulting polymerase ribozyme, termed tC19Z, can synthesize a functional copy of a small hammerhead ribozyme, the first example of transcription of a ribozyme by a ribozyme. However, the polymerase remained strongly biased toward unstructured, C-rich templates,^12^ which prevents the synthesis of more complex products.

Subsequent efforts to evolve the polymerase ribozyme required it to synthesize progressively larger functional RNAs, with selection of the ribozymes based on the activity of an attached RNA product they had synthesized. This strategy links polymerase genotype to product phenotype, thereby providing selection pressure for the efficient, general, and accurate synthesis of RNA. The initial functional products to be synthesized were portions of two different RNA aptamers, requiring the addition of 18–30 nucleotides.^7^ Following 24 rounds of this procedure, the 24-3 polymerase was obtained, which has ∼100-fold faster catalytic rate compared to prior polymerases and can synthesize a complete, functional 49-nucleotide ribozyme.

Next, polymerases were selected based on their ability to synthesize a complete hammerhead ribozyme, which yielded the 38-6 polymerase after 14 additional rounds of evolution,^13^ then the 52-2 polymerase after 14 more rounds.^14^ By that point, polymerase activity had improved by more than 600-fold and the polymerase gained the ability to bind the primer-template duplex with 1000-fold enhanced affinity.^15^ Sequence generality was also substantially improved. Although highly structured, mixed-sequence RNA templates of >50 nucleotides remain difficult for the 52-2 polymerase to copy in high yield, it is able to synthesize its own ancestor, a functional class I ligase.

Early polymerase ribozymes could operate with 96–99% fidelity on the most favorable template sequences, but were unable to operate on difficult sequences.^6,11^ The enhanced generality of the improved polymerases came at a cost to fidelity, which was reduced to 92% per nucleotide for typical templates, and even lower for longer and more difficult templates.^7,13^ The polymerase ribozyme tends to stall or terminate extension after a mismatched nucleotide has been incorporated. In early variants, high rates of termination led to poor yield, even for favorable sequences, with the truncated products having a high level of 3’-terminal mutations.^12^ For the improved variants, 24-3, 38-6, and 52-2, these terminal mutations could be extended, albeit at lower rates, leading to an accumulation of more mutated full-length RNA products.^13^ The challenge was particularly acute for synthesis of complex RNAs, with the 52-2 polymerase synthesizing the class I ligase with an error rate of 16% per nucleotide, resulting in only 0.1% of the full-length RNA products able to function as a ligase.^14^

The ability of the improved polymerases to synthesize the class I ligase provided a lever to improve fidelity. The population of polymerase ribozymes from which 52-2 was isolated were subjected to selection for synthesis of a functional class I ligase, imposing strong selection pressure to favor more accurate polymerases. To further enhance fidelity, the concentration of the catalytic Mg^2+^ cofactor was reduced over the course of evolution, from 200 to 83 mM. In previously reported work, polymerase variant 71-89, isolated after the 19th round of this period of directed evolution (71 total rounds involving synthesis of functional RNAs), showed both enhanced activity and improved fidelity in the presence of 50 mM Mg^2+^.^16^ This improvement enabled the 71-89 polymerase to replicate the hammerhead ribozyme, synthesizing both the ribozyme and its complement at 50 mM Mg^2+^ with an average fidelity of 94.4% per nucleotide. In contrast, the 52-2 polymerase requires 200 mM Mg^2+^ to synthesize the hammerhead ribozyme, and does so with only 90.2% fidelity. This difference placed the two polymerases on either side of the error threshold, with hammerhead ribozymes replicated by 52-2 decaying into nonsense sequences, while those replicated by 71-89 retaining function and undergoing adaptive evolution.

As described in the present study, directed evolution based on synthesis of the class I ligase was continued for an additional 14 rounds, while further reducing the concentration of Mg^2+^ to 50 mM. Sequence analysis of the evolving population revealed complex dynamics involving competing clades of polymerases with distinct adaptive responses to the selection constraints. Two clades that were prevalent in the final round have an improved ability to synthesize the class I ligase at reduced Mg^2+^ concentrations. Combining mutational features from these two clades yielded the 85h34 polymerase, which differs from 52-2 by 20 mutations.

The 85h34 polymerase can synthesize a diverse repertoire of functional RNAs in high yield under reduced Mg^2+^ concentrations, including aptamers and endonuclease and ligase ribozymes. The 85h34 polymerase retains activity even at 10 mM Mg^2+^, which enables it to achieve a fidelity of 94–95% per-nucleotide across multiple functional RNA targets, including the class I ligase. The enhanced sequence generality and fidelity of 85h34 demonstrates the broad capacity of an RNA polymerase ribozyme to support the transcription and expression of a variety of functional RNA elements, with direct relevance to the RNA world hypothesis. Furthermore, the RNA-catalyzed evolution of other functional RNAs, first demonstrated with 71-89, may now be expanded to larger and more complex RNAs.

## Materials and Methods

### Materials

The sequences of all oligonucleotides used in this study are provided in Table S1. Synthetic oligonucleotides were either purchased from IDT or prepared by solid-phase synthesis using an Expedite 8909 DNA/RNA synthesizer, with reagents and phosphoramidites from either Glen Research or ChemGenes. All polymerase ribozymes and templates for RNA-catalyzed RNA synthesis were prepared by *in vitro* transcription from corresponding DNA templates. All RNA products were purified by denaturing polyacrylamide gel electrophoresis (PAGE) and quantified by UV absorbance on a Nanodrop 2000 spectrophotometer. Primer extension assays with polymerase ribozymes used fluorescently-labeled RNA primers, and the products were separated by PAGE, imaged using an Amersham Typhoon RGB laser scanner, quantified using ImageQuant TL software, and the data plotted using GraphPad Prism. Hot Start Taq, Q5 high-fidelity DNA polymerase, T4 RNA ligase, K227Q T4 RNA Ligase 2, and Universal miRNA Cloning Linker were from New England Biolabs. SuperScript IV reverse transcriptase, Turbo DNase, and MyOne C1 streptavidin magnetic beads were from Thermo Fisher Scientific. Nucleoside 5’-triphosphates, pCp-biotin, and γ-(2 azidoethyl)-ATP were from Jena Bioscience, and all other chemical reagents were from Sigma-Aldrich.

### *In vitro* transcription

To prepare the successive populations of polymerase ribozymes, the T7 RNA polymerase promoter was installed during the PCR amplification step of evolution, allowing the resulting DNAs to be used directly for *in vitro* transcription. To prepare individual polymerases, corresponding DNA templates were obtained by PCR amplification of a portion of a plasmid that encodes the polymerase, which was purchased from IDT as a “Rapid Gene”. To prepare RNA templates for RNA-catalyzed polymerization, corresponding DNA templates were obtained either by PCR assembly or cross-extension of oligodeoxynucleotides. PCR assembly used 1 nM input DNA, 0.5 μM primers, and Hot Start Q5 DNA polymerase according to the manufacturer’s protocol, heating at 98 °C for 90 s, then 25 cycles of 98 °C for 10 s, 52 °C for 10 s, and 72 °C for 30 s, and final heating at 72 °C for 5 min. DNA cross-extension used 5 µM oligodeoxynucleotides and Superscript IV according to the manufacturer’s protocol. All DNA products were purified using the New England Biolabs Monarch PCR and DNA cleanup kit.

*In vitro* transcription was carried out in a mixture containing 20 ng/μL DNA, 15 U/μL T7 RNA polymerase, 0.002 U/μL inorganic pyrophosphatase, 5 mM each NTP, 25 mM MgCl_2_, 2 mM spermidine, 10 mM DTT, and 40 mM Tris-HCl (pH 8.0), which were incubated at 37 °C for 2 h, followed by addition of 0.1 U/μL Turbo DNase and incubation at 37 °C for 1 h. The reactions were quenched by adding excess EDTA relative to MgCl_2_ and the products were purified by PAGE.

### *In vitro* evolution

*In vitro* evolution of polymerase ribozymes based on the ability to synthesize a functional class I ligase was carried out as previously described.^16^ The starting population, obtained after the prior 52 rounds of evolution,^14^ was subjected to error-prone PCR,^17^ and a new 5’-terminal region containing an AvaII restriction site was installed by PCR using primers Fwd1 and Rev1. *In vitro* transcription of polymerase RNAs was carried out in the presence of γ-(2-azidoethyl)-ATP to enable attachment via click chemistry of a 5’-hexynylated RNA primer (Primer1). Following PAGE purification, 50 nM polymerase-primer conjugates were mixed with 100 nM 3’-biotinylated RNA template (Tem2) and 75 nM cofactor oligodeoxynucleotide (Tem4), which were annealed by heating at 80 °C for 30 s, then cooling to 17 °C. The cofactor oligonucleotide binds to the new 5’-terminal region and was found to increase polymerase activity.

The annealed materials were added to a reaction mixture containing the four NTPs, various concentrations of MgCl_2_, 50 mM Tris-HCl (pH 8.3), and 0.05% Tween-20, which were incubated at 25 °C for various times. The MgCl_2_ and NTP concentrations and reaction times over the course of evolution are listed in Table S2. The reaction was quenched with EDTA and mixed with streptavidin-coated magnetic beads to capture the biotinylated templates. The beads were washed twice with urea buffer (8 M urea, 1 mM EDTA, and 10 mM Tris-HCl (pH 8.0)), and the polymerase-primer conjugates were eluted from the RNA template in NaOH buffer (25 mM NaOH, 1 mM EDTA, and 0.05% Tween-20), then neutralized in 100 mM Tris HCl (pH 7.5), ethanol precipitated, and the products were purified by PAGE.

The polymerases with attached RNA product were captured on magnetic beads that had been derivatized with an oligodeoxynucleotide (Rev2) that is complementary to the 3’ end of the polymerase, then reverse transcribed *in situ*. In a concerted reaction using both AvaII type IIs restriction enzyme and T4 DNA ligase, the 5’ substrate for the class I ligase (S1) was installed at the 3’ end of the polymerase cDNA. The 3’ substrate (S2), which was 3’-biotinylated, was added to the solution and the bead-bound ligase ribozymes were challenged to join the two substrates in the presence of 60 mM MgCl_2_, 200 mM KCl, 0.6 mM EDTA, and 50 mM Tris-HCl (pH 8.3) at 25 °C for 30 min.

All RNAs not attached to cDNA were washed from the beads using NaOH buffer, then the polymerase cDNAs and any ligated substrates were cleaved off the beads using EcoRV, which cleaves at a site within the 5’ primer region of the cDNA, assisted by a complementary oligodeoxynucleotide (Tem8). The released molecules were captured on streptavidin-coated magnetic beads, washed three times with NaOH buffer and twice with urea buffer, eluted in formamide buffer (95% formamide, 100 nM EDTA, and 1 mM Tris-HCl (pH 8.0)) by heating at 95 °C for 5 min, then ethanol precipitated. The cDNAs were reverse transcribed using primer Fwd2, which is specific for substrate S2, then PCR amplified using primers Rev1 and Fwd3, the latter of which is specific for the ligation junction between S1 and S2. Nested PCR using primers Fwd4 and Rev1 provided materials to begin the next round of evolution. New mutations were introduced by error-prone PCR at various rounds, as shown in Table S2.

### Analysis of the evolving population by deep sequencing

Illumina adapter and index sequences were added to PCR products obtained after various rounds of evolution by nested PCR with primers Fwd5 and Rev6 and the Illumina Nextera Index primers. Sequencing was performed by the Salk Institute Next Generation Sequencing Core on an Illumina NextSeq 2000 with a 150-cycle paired end run, which generated ∼5 million paired reads per sample. The adapter and primer sequences were trimmed using cutadapt v2.4,^18^ the paired reads were merged using the paired-end read merger program PEAR (v 0.9.11),^19^ and the merged reads were filtered to recover only those with a quality score ≥30 at every position using FASTX Toolkit v0.0.14.^20^ Using a custom Python script, the occurrence of each unique sequence was tallied across all sequenced rounds, including rounds 43–52 that had been sequenced in a previous study.^14^ Unique sequences were aligned to the ancestral 38-6 polymerase sequence,^13^ and the frequency of each distinct mutation relative to the 38-6 polymerase was enumerated across sequences and rounds. Using only mutations that reached a prevalence of at least 0.1% in at least one round, unique sequences were converted to a binary presence-or-absence vector, and these high-dimensional mutation vectors were embedded into a three-dimensional representation using the PaCMAP dimension-reduction algorithm.^21^ Genotypes were grouped into hierarchical clusters based on proximity in the embedded space and shared representation over the same rounds, and a phylogenetic tree was built from these clusters by maximum parsimony. The three-dimensional projection of the genotypes, colored by clade membership, and a Muller plot of clade frequency over the rounds were generated using Plotly.^22^

### RNA-catalyzed polymerization of RNA

RNA polymerization reactions were conducted as described previously,^14^ but with a polymerase folding step prior to initiating the reaction. 100 nM polymerase ribozyme, 80 nM Cy5- or fluorescein-labeled and 3’-biotinylated RNA primer, and 100 nM RNA template were mixed and annealed by heating at 80 °C for 30 s and then cooling to 17 °C. The annealed RNAs were added to a reaction mixture containing 10, 50, or 200 mM MgCl_2_, 50 mM Tris-HCl (pH 8.3), and 0.05% Tween-20, with 8% (w/v) PEG8000 added when using 10 mM MgCl_2_. The mixture was incubated for 1 h at the specified temperature, typically 25 °C for 200 and 50 mM MgCl_2_ and 27 °C for 10 mM MgCl_2_, which allowed the polymerase to fold. To initiate the reaction, an equal volume of a solution containing the same components as above plus the four NTPs was added, mixed, and incubated at the same temperature. The final NTP concentrations were 0.5, 2, or 4 mM for 10, 50, or 200 mM MgCl_2_, respectively. The reaction was quenched by adding an equal volume of a solution containing 250 mM EDTA (pH 8.0), 500 mM NaCl, 5 mM Tris-HCl (pH 8.0), and 0.025% Tween-20, then mixed with 5 µg streptavidin-coated magnetic beads per pmol biotinylated RNA primer, and incubated with gentle agitation at 25 °C for 30 min. Prior to use, the beads had been washed according to the manufacturer’s protocol, then incubated with a solution containing 1 mg/mL tRNA, 1 M NaCl, 1 mM EDTA, and 10 mM Tris-HCl (pH 8.0) for 30 min. The RNA template was removed from the bead-bound materials by two washes with NaOH buffer, followed by one wash with urea buffer. Then the reaction products were eluted from the beads by incubating in 95% formamide and 10 mM EDTA at 95 °C for 10 min, and were analyzed by PAGE. For reactions of the 85h34 polymerase with the short, repeat, and hairpin RNA templates, a version of the ribozyme with the original pairing to the template RNA was used (85h34.1), while for the synthesis of functional RNAs a version with more stable pairing to the template was used (85h34.2).

### Analysis of polymerase fidelity

The polymerase ribozyme synthesized the class I ligase as described above. The eluted materials were purified by PAGE to obtain the full-length products, which were converted to corresponding DNA libraries as previously described,^14^ and sequenced by the Salk Institute Next Generation Sequencing Core on an Illumina MiniSeq with a 150-cycle paired-end run.

For analysis of the synthesized hammerhead, F1 ligase, and class I ligase, including both partial and full-length products, after the materials were captured on streptavidin-coated beads and washed with NaOH and urea, as described above, the beads were washed twice with TE buffer (1 mM EDTA and 10 mM Tris-HCl (pH 8.0)) and once with New England Biolabs T4 RNA Ligase buffer supplemented with 0.05% Tween-20. The bead-bound RNA products were ligated to the Universal miRNA Cloning Linker using K227Q T4 RNA Ligase 2 according to the manufacturer’s protocol, but adding 20% PEG8000 and 0.05% Tween-20. After incubating at 23 °C for 4 h with gentle agitation, the beads were washed twice with urea buffer and twice with TES buffer (1 M NaCl, 1 mM EDTA, and 10 mM Tris-HCl (pH 8.0)), then incubated for 15 min with 1 μM primer Rev611 in TES buffer with gentle agitation. The RNA was then reverse transcribed *in situ* at 50°C for 1 h using primer Rev7 and Superscript IV according to the manufacturer’s protocol, but adding 0.05% Tween-20. Finally, the beads were washed twice with TE buffer and the cDNAs were eluted in two washes of NaOH buffer, which was neutralized in 100 mM Tris HCl (pH 7.5). The recovered cDNAs were converted to double-stranded DNA by second-strand synthesis using primers Fwd8, Fwd9, Fwd10, and Fwd11 for Mango aptamer, hammerhead, F1 ligase, and class I ligase products, respectively, then prepared for sequencing by PCR using Q5 DNA polymerase according to the manufacturer’s protocol. During the first 5 cycles of PCR Illumina adapters were added (Fwd12 and Rev8), and in the next 8 cycles indexes were added using the Illumina Nextera Index primers. The final double-stranded DNAs, containing 125–210 nucleotides, were sequenced by the Salk Institute Next Generation Sequencing Core on an AVITI Element instrument with a 150-cycle paired-end run.

The sequencing data were processed using a custom bioinformatics pipeline to categorize all mutations relative to the expected sequence, separately tabulated for each size of partial- and full-length products. The adapter and primer sequences were trimmed using cutadapt v2.4,^18^ the paired reads were merged using FLASH v1.2.11,^23^ and the merged reads were filtered to recover only those with a quality score ≥40 at every position using FASTX Toolkit v0.0.14.^20^ A Python script was used to calculate the numbers of matches, mismatches, deletions, and insertions as a function of template position and read length along the reference sequence. The resulting data tables were processed manually to obtain fidelity values and position-specific data plots.

## Results

### Evolution of Polymerases with Improved Activity and Fidelity

Fifty-two prior rounds of directed evolution, each requiring the polymerase ribozyme to synthesize a functional RNA, resulted in the 52-2 polymerase ribozyme, which has markedly improved activity compared to its predecessors.^14^ The 52-2 polymerase has the ability to synthesize complex structured RNAs, including its own ancestor the class I ligase. However, the polymerase exhibits poor fidelity when synthesizing such complex RNAs molecules, with an average fidelity of only 84% per nucleotide when synthesizing the class I ligase. During the 52 rounds of evolution, the polymerase was required to synthesize small functional RNAs, including aptamers and the hammerhead ribozyme, which contain only 10–15 critical nucleotide positions that must be copied accurately to yield a functional product. Thus there was only modest selection pressure during those rounds for accurate synthesis. In contrast, the class I ligase contains 97 nucleotides, more than half of which are critical for function.^10^ Accordingly, the poor fidelity of the 52-2 polymerase results in class I ligase products that are more than 1000-fold less active compared to those produced by *in vitro* transcription using T7 RNA polymerase protein. It was reasoned that further evolution of the polymerase ribozyme based on its ability to synthesize a functional class I ligase ribozymes would impose strong selection pressure on copying fidelity, while also requiring the synthesis of a large and complex RNA.

To select for RNA-catalyzed synthesis of the class I ligase, the population of polymerase ribozymes were joined to an RNA primer via a 5’,5’-linked flexible tether (Figure 1a). The evolving polymerases were required to extend the attached primer on a separate RNA template that encodes the class I ligase. The resulting products were captured by hybridization of the 3’ end of the polymerase to oligodeoxynucleotide-coated beads, then the polymerases were reverse transcribed to render them inactive as an RNA:cDNA duplex, with the RNA strand attached to the primer-extension product. One substrate for the ligase, bearing a free 3’-hydroxyl, was then attached to the 3’ end of the cDNA. The other substrate, bearing a 5’-triphosphosphate, was provided as a separate molecule, connected via a short hairpin to the template for ligation, and with a biotin moiety at the 3’ end. Only those polymerase ribozymes that had successfully synthesized a catalytically active ligase were able to join the two substrates, thus linking the 3’-biotin to the polymerase-encoding cDNA. The cDNAs were released from the oligodeoxynucleotide-coated beads by site-specific DNA cleavage, then the biotinylated products were captured on streptavidin-coated beads and the cDNAs were selectively amplified by PCR and forward transcribed to generate progeny polymerases.

**Figure 1.**
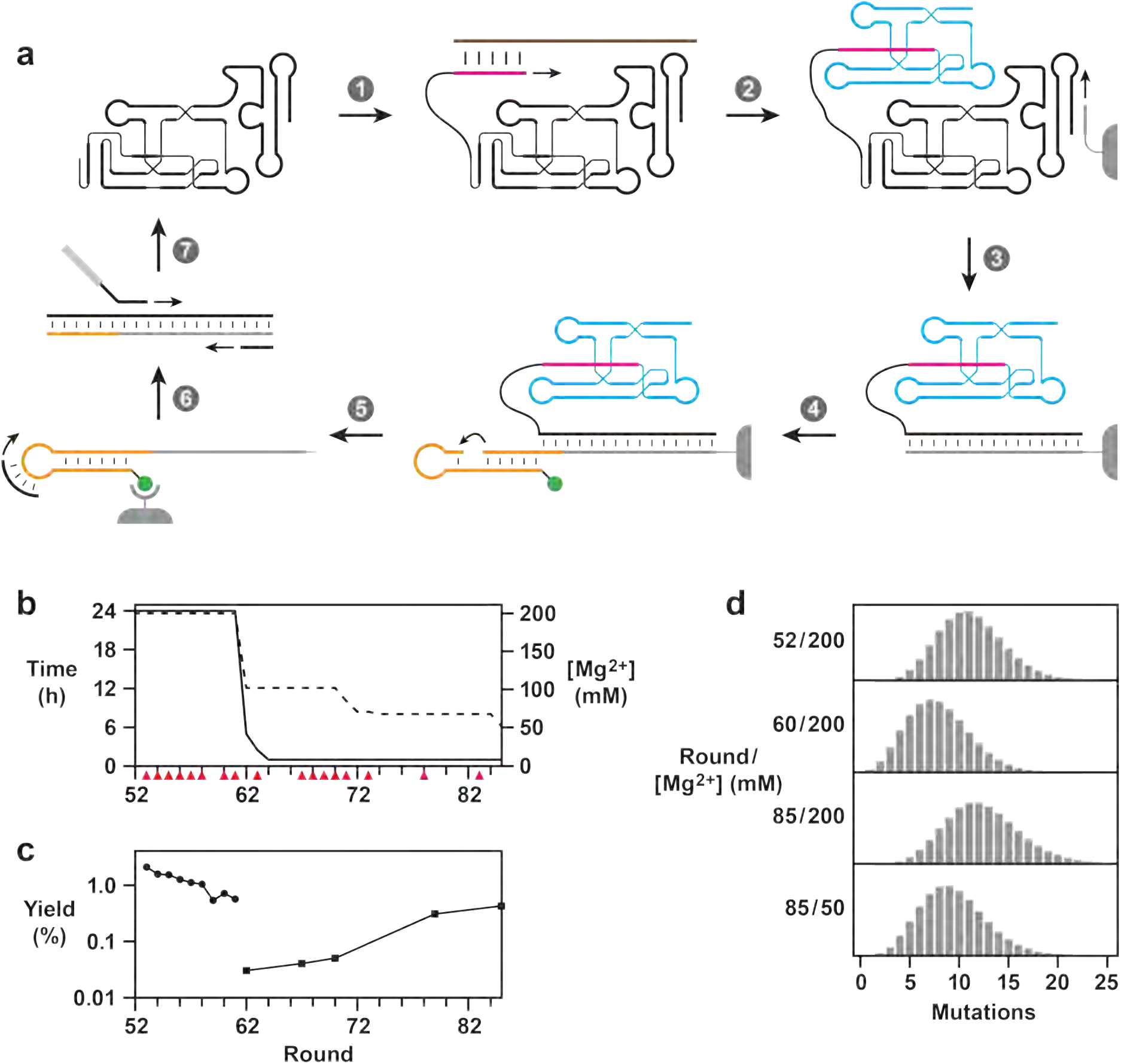
Directed evolution of the 85h34 polymerase. (a) Scheme for selective amplification of polymerase ribozymes that synthesize a functional class I ligase (blue). (*1*) Attachment of an RNA primer (magenta) that binds to an RNA template (brown) encoding the ligase; (*2*) extension of the primer to form the ligase; (*3*) reverse transcription of the polymerase using a bead-bound DNA primer (gray); (*4*) attachment to the cDNA of one of the substrates for ligation (orange), which binds to a template that is linked to the other substrate and is biotinylated (green circle); (*5*) release of the ligated products from the bead and subsequent capture on streptavidin; (*6*,*7*) selective PCR amplification and forward transcription to generate progeny polymerases. (b) Reaction time and Mg^2+^ concentration for polymerization and use of mutagenic PCR (red triangles) during rounds 53–85. (c) Yield after either 24 h (rounds 52–61) or 1 h (rounds 62–85) for synthesis of the class I ligase by the population of polymerases in the presence of 200 mM MgCl_2_ and 4 mM NTPs at pH 8.3 and 25 °C. (d) Distribution of mutations in class I ligase ribozymes that were synthesized in 24 h by the population of polymerases obtained after rounds 52, 60, and 85. Reaction conditions were as above, except for the round 85 population at 50 mM MgCl_2_, where the concentration of NTPs was 2 mM.

The first round of this demanding evolution procedure was initiated using the entire population of polymerase ribozymes resulting from the prior 52 rounds, further mutagenized by error-prone PCR.^17^ Thirty-three additional rounds of evolution were carried out (rounds 53–85), each requiring the RNA-catalyzed synthesis of a functional class I ligase. The stringency of selection was increased over the course of evolution in two ways. First, the reaction time for polymerization was reduced from 24 h (rounds 53–61) to 1 h (rounds 64–85). Second, the concentration of Mg^2+^ was reduced from 200 mM (rounds 53–61) to 100 mM (rounds 62–70) and ultimately to 50 mM by round 85 (Figure 1b; Table S2). Reducing the Mg^2+^ concentration places increased demands on RNA catalysis, while destabilizing base mismatches, especially G:U wobble pairs, to provide indirect selection pressure for improved fidelity.^12^ Further mutagenesis by error-prone PCR was carried out intermittingly during the course of evolution to maintain genetic diversity in the population.

The response of the evolving population to these various selection pressures points to multiple distinct adaptations that improved the ability of the polymerase to synthesize a functional ligase. During rounds 53–61, the fidelity of synthesis (in the presence of 200 mM Mg^2+^) improved from 86% to 91% per nucleotide, but the yield of ligase products decreased by nearly 4-fold (Figure 1c). Nonetheless, the specific activity of these products increased by 10-fold due to the significantly lower frequency of mutations (Figure S1; Figure 1d). Beginning at round 62, when the concentration of Mg^2+^ was reduced from 200 to 100 mM, and following round 64, when the reaction time was reduced from 24 to 1h, there was a marked improvement in polymerase activity. This improvement continued after round 70 when the concentration of Mg^2+^ was further reduced. Compared to the population of ribozymes at round 52, the population at round 85 exhibited improved yield of the class I ligase in the presence of either 200 or 50 mM Mg^2+^, similar fidelity at 200 mM Mg^2+^ (85.6% vs. 86.3%), and improved fidelity at 50 mM Mg^2+^ (89.5%), a condition where the 52-2 population is unable to synthesize full-length products.

### Population dynamics over the course of evolution

Deep sequencing was carried out for each of the polymerase populations from rounds 53 through 85 to reveal the progression of acquired mutations relative to the 52-2 polymerase. Although many mutations quickly swept to fixation, some displayed more complex dynamics, rising early and then collapsing, or persisting at a low level in the population. There are several correlated pairs of mutations that rose and fell together, located at paired positions within stem regions of the polymerase, including P4, P6, P7, and P8 of the catalytic core domain and A4 of the accessory domain (Figure 2a). Broader clusters of mutations rose and fell in concert and appear to have occurred in three distinct epochs (rounds 58–62, 67–75, and 79–85) that correlate with notable changes in the stringency of selection (Figure 1b).

**Figure 2.**
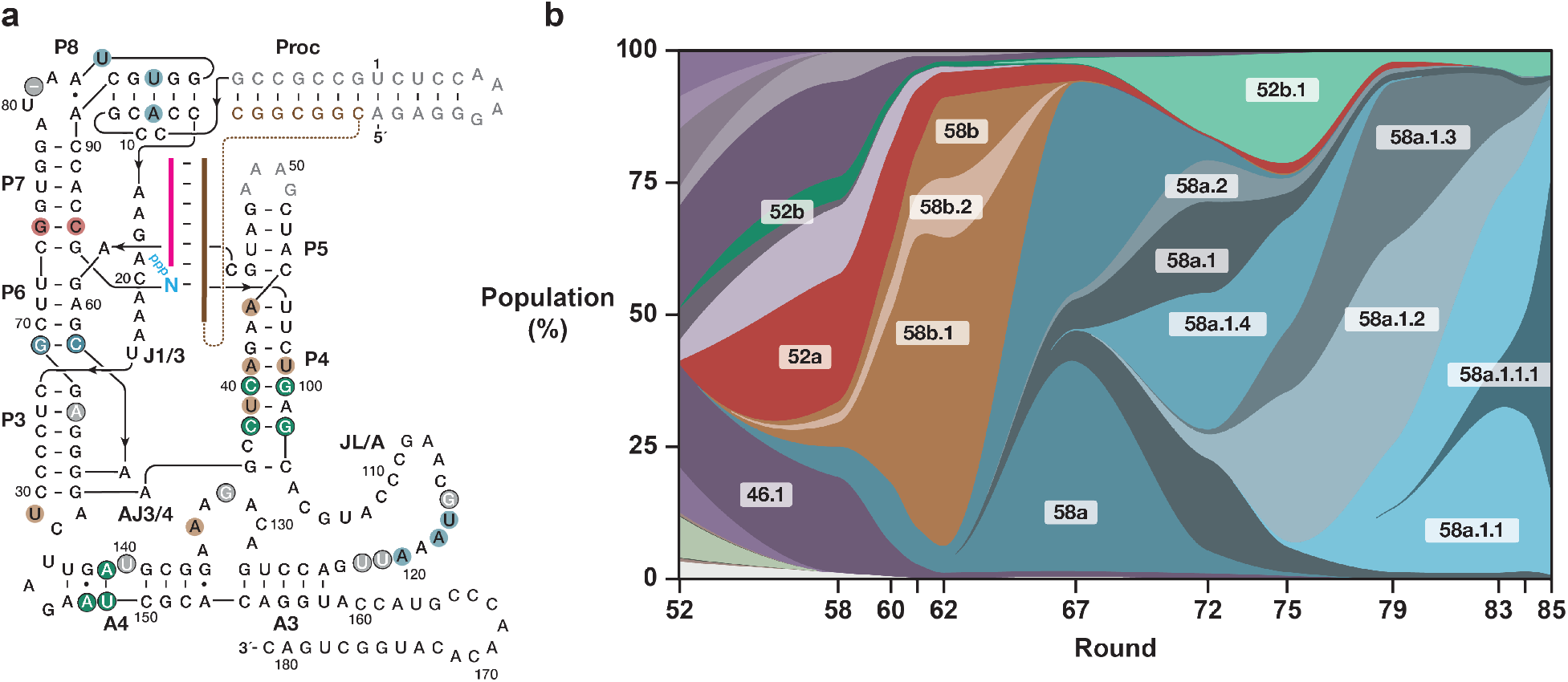
Accumulation of mutations over the course of evolution. (a) Mapping of mutations onto the secondary structure of the polymerase. Paired regions are labeled, including the processivity tag pairing between ribozyme and template (Proc). Mutations present in the 85h34 polymerase relative to 52-2 are highlighted with circled white letters, with the color of the circle corresponding to the clade in which they arose (or gray for mutations that arose in multiple clades). Other mutations that were present in major clades but not in 85h34 are highlighted with circled black letters, with the circles colored similarly. (b) Muller plot of evolving clades over successive rounds of evolution, with the area corresponding to population frequency of the clade over time. Mutations in (a) and regions in (b) are colored similarly based on clade membership. The 52-2 clade and other descendants of the ancestral clade from round 46 are shown in purple. The major successor clades are 52a (red), 58b (brown), 58a (blue), and 52b (green). Sub-clades of these that reached a frequency of ≥5% of the population are shown as distinct bands with different shades of the parent color.

To better visualize the population dynamics, the distribution of polymerase sequences for rounds 39 to 85 was projected to three dimensions using a dimension reduction algorithm that attempts to maintain relative short- and long-range distances between datapoints. The prior rounds of evolution involving synthesis of the hammerhead ribozyme (rounds 39–52) were included in the analysis to incorporate population dynamics leading up to the period of evolution that required synthesis of the class I ligase. The resulting three-dimensional embedding of the evolving sequence landscape shows dense clusters joined by sparse filaments (Figure S2a). Individual clusters tend to be populated over a narrow interval of successive rounds, each representing diverse collections of sequences that have similar trajectories of population frequency and share a common set of linked mutations. Filaments appear to be transitional forms between clusters, adding, removing, or blending linked sets of mutations between distinct clusters. Viewed over time, population density occupies clusters in succession, often transiently occupying the filaments as the population shifts to new clusters. Overall, the pattern is consistent with quasispecies evolution,^8^ with clouds of related sequences exploring a dynamic fitness landscape, as would be expected from the high rates of mutagenesis and stringency of selection employed throughout the evolution procedure.

A custom hierarchical clustering algorithm, using both group proximity in the embedded three-dimensional space and a maximum parsimony phylogenetic reconstruction principle, ordered the disparate clusters into a phylogenetic tree, revealing broad quasispecies clade dynamics across the evolutionary lineage (Figure S2b). At round 52 there is a dense network of closely related clusters, all sharing features of the dominant 52-2 sequence. Beginning at round 58 there was an explosion of new clusters, all derived from 52-2, adding new mutational features to the shared 52-2 core sequence. However, only four such clusters gave rise to persistent clades of linked clusters that share common features and persisted over many subsequent rounds, both increasing in abundance and accumulating additional mutations. A Muller plot of the representation of these clades in the population shows the complex pattern of succession that occurred over rounds 52–85 (Figure 2b). Two such clades peaked early in the first epoch of evolution, before collapsing. Clade 52a emerged before selection for ligase synthesis had begun, detectable as a satellite cluster to the 52-2 cluster in round 52 and earlier, peaking at 20% of the population in round 58, declining to 5% of the population by round 62, and disappearing entirely by round 79. This clade is defined by replacement of an A:U pair by G:C within the P7 stem, although additional mutations in the P3 and A4 regions that later become prevalent in other clades are first seen in this clade (Figure S2b). Clade 58b arose later, first appearing in round 58, reaching dominance in the population at round 61, then collapsing to extinction after the second epoch began when selection stringency was increased (Figure 1b). This clade is defined by a set of mutations in the P4 stem that weaken pairing near the bulged C37 nucleotide that helps form the active site of the polymerase, and contains a dominant subclade (58b.1) with an A-to-U mutation in the AJ3/4 loop that replaces one of the adenosines that participates in the “adenine stack”, a critical tertiary interaction that scaffolds the nucleotide binding pocket.^24^ Consistent with these weakened tertiary contacts in the active site and NTP binding pocket, the rise of clade 58b closely correlates with a period when polymerase activity decreased and fidelity increased.

Two clades that were present at low levels in the early rounds came to displace the early dominant clades as selection pressure was increased. Clade 58a is initially defined by covarying mutations in the P6 stem that convert the G62:C69 pair to C:G. This clade persisted at 5–10% frequency during rounds 58–62 before rising to >90% of the population as clade 58b disappeared. Subclade 58a.1 displaced 58a by round 72 after acquiring a second covarying pair in P8 that converts A13:U86 to U:A, together with mutation of the nearby unpaired U83 to C. Clade 52b arose in early rounds and persisted at <5% frequency in parallel with clade 58a, initially defined by covarying mutations in P4 that convert G40:C100 to C:G. Following round 62, subclade 52b.1 rose to >20% of the population by round 75, acquiring additional covarying mutations in P4 that converted G38:C102 to C:G and in A4 of the accessory domain that converted U141:A149 to A:U. Together, the 58a and 52b clades account for 95–99% of all sequences from round 72 onward. These two clades competed for population share, with 58a first declining as 52b emerged, then becoming dominant again as 52b declined.

Although the distinguishing mutations in clades 58a and 52b remain separate, the two clades also accumulate common mutational features. In both clades, G67 was mutated to A, converting a G:U wobble pair in the P3 stem to A:U; C80 of the J7/8 loop was mutated to U and later to a deletion; A132 of the AJ3/4 loop was mutated to G; and bulged nucleotide A140 in the A4 region was mutated to U. Following the second epoch of evolution, from round 75 onward, both clades sampled various mutations in the distal portion of the JL/A linker, which joins the polymerase core and accessory domains. Ultimately, each clade settled on a distinct sequence in this region.

Recombination appears to have been prevalent between the 58a and 52b clades in later rounds of evolution. Most such events appear to be non-productive, resulting from crossover within the core of the polymerase that combine only one member of a covarying pair from P8 of 58a and P4 of 52b, causing base mismatches in both stems. However, crossover in the JL/A linker resulted in mutations in the A4 region of the accessory domain that first appeared in 52b and later appeared in the 58a.1.1.1 subclade of 58a, becoming dominant in the population by round 85.

### An Improved Hybrid Polymerase

The long persistence of two clades with distinct sets of mutations could represent mutually incompatible adaptations, but may also reflect clonal interference. The latter seems plausible given that both clades are defined by paired mutations that are well separated in the primary sequence, preventing the transfer of the paired mutations through a single recombination event. In support of this hypothesis, an engineered variant that combines the C62:G69 mutations in P6 of clade 58a with the C38:G102 mutations in P4 of clade 52b exhibited improved activity in synthesizing the class I ligase, especially in the presence of 50 mM Mg^2+^. This variant was further modified by replacing the 5’-terminal sequence used during evolution by a simple 5’ hairpin. The resulting polymerase was named “85h34”, in consideration of it being the 34th hybrid tested, derived from the 85th round of evolution.

The 85h34 polymerase ribozyme has dramatically improved ability to synthesize the class I ligase compared to the 52-2 polymerase, as well as the 71-89 polymerase that was isolated following round 71 and previously shown to have improved activity and fidelity in the presence of 50 mM Mg^2+^.^16^ 71-89 shares some, but not all, of the mutations characteristic of the 52b clade, including C38:G102 in P4 and G132 in AJ3/4. RNA-catalyzed synthesis of the class I ligase by the 52-2, 71-89, and 85h34 polymerases was tested in the presence of 200, 50, and 10 mM Mg^2+^ (Figure 3). At 200 mM Mg^2+^, all three polymerases yielded full-length products after 24 h, although the 71-89 and 85h34 polymerases gave >10-fold higher yield compared to 52-2. At 50 mM Mg^2+^, 52-2 can no longer synthesize the full-length ligase, 71-89 has reduced yield, and 85h34 maintains high yield. At 10 mM Mg^2+^, only 85h34 retains substantial activity. Sequencing the full-length products synthesized by the 85h34 polymerase in the presence of 50 mM Mg^2+^ revealed an average fidelity of 92.3% per nucleotide, exceeding that of any other polymerase tested, including the 52-2 polymerase at 200 mM Mg^2+^.

**Figure 3.**
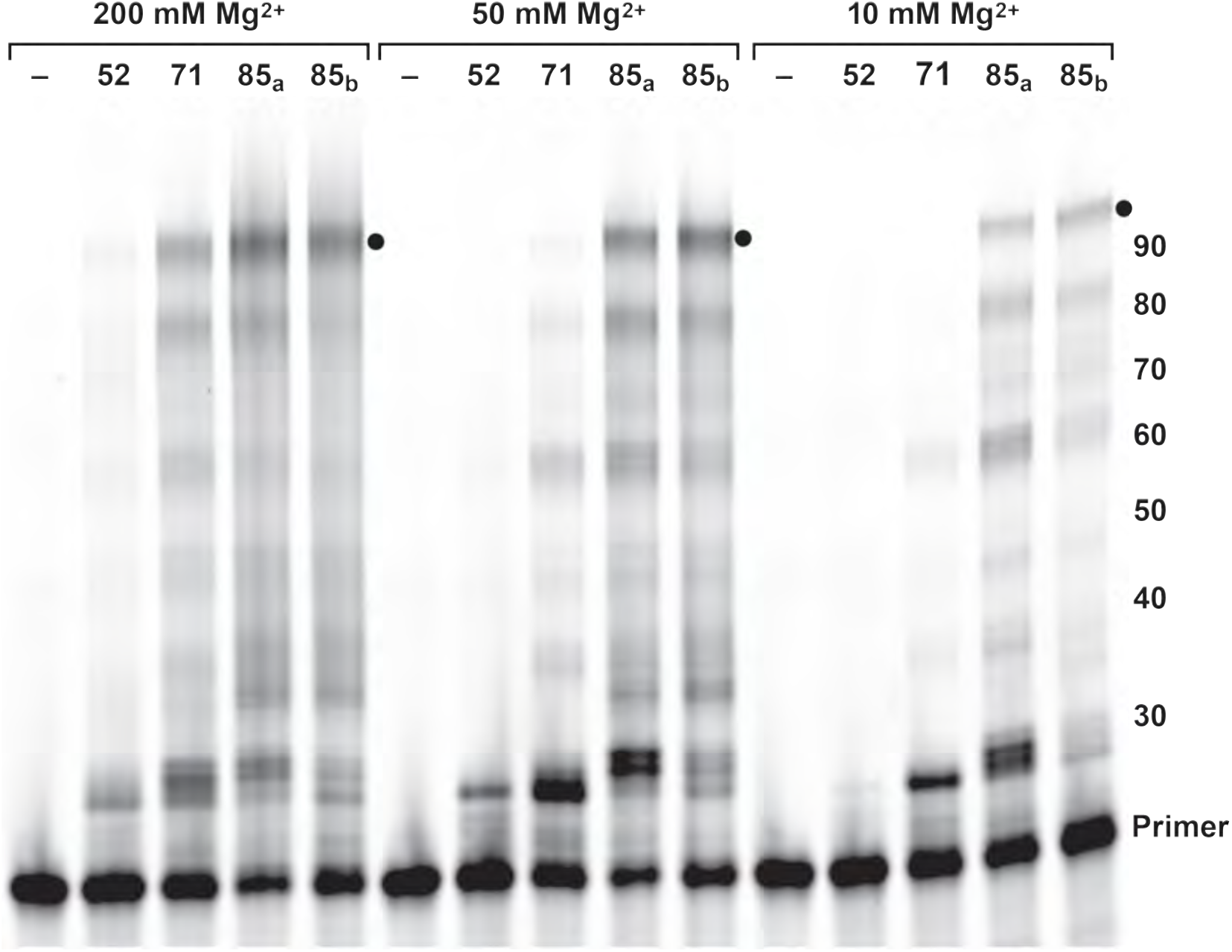
RNA-catalyzed synthesis of the class I ligase. The 52-2, 71-89, and 85h34 polymerases, obtained after rounds 52, 71, and 85, respectively, were tested for the ability to synthesize the class I ligase in the presence of 200, 50, or 10 mM Mg^2+^. Two different forms of 85h34 were tested (85a and 85b) that differ with regard to the sequence of the processivity tag that binds the 5’ end of the template. The concentration of NTPs was 4, 2, or 0.5 mM for reactions carried out in the presence of 200, 50, or 10 mM Mg^2+^, respectively. All reactions were at pH 8.3 and 25 °C for 24 h and the products were separated by PAGE. Black dots indicate full-length products.

### Copying of Various RNA Sequences and Structures

Early polymerases in the lineage derived from the class I ligase were limited to copying unstructured, C-rich templates. To evaluate the capacity of 85h34 to operate on a variety of template sequences, four templates of 11–12 nucleotides were compared: **t1**, which has long been used as a standard template to evaluate polymerase extension rates under the most favorable conditions, and **tA**, **tB**, and **tC**, each chosen as an arbitrary sequence with balanced nucleotide composition. Notably, **t1** contains only a single instance of two consecutive nucleotides that do not include cytosine (the central UU), while **tA**, **tB**, and **tC** contain 6–8 such consecutive pairs. Although both 52-2 and 85h34 yield full-length copies of **t1** within 15 s (Figure 4a), 52-2 struggles to complete copying of the other three templates, stalling at template regions of two or more consecutive purines. 85h34 also slows at these regions, but completes the synthesis within 30–100 s for all three of the more challenging templates.

**Figure 4.**
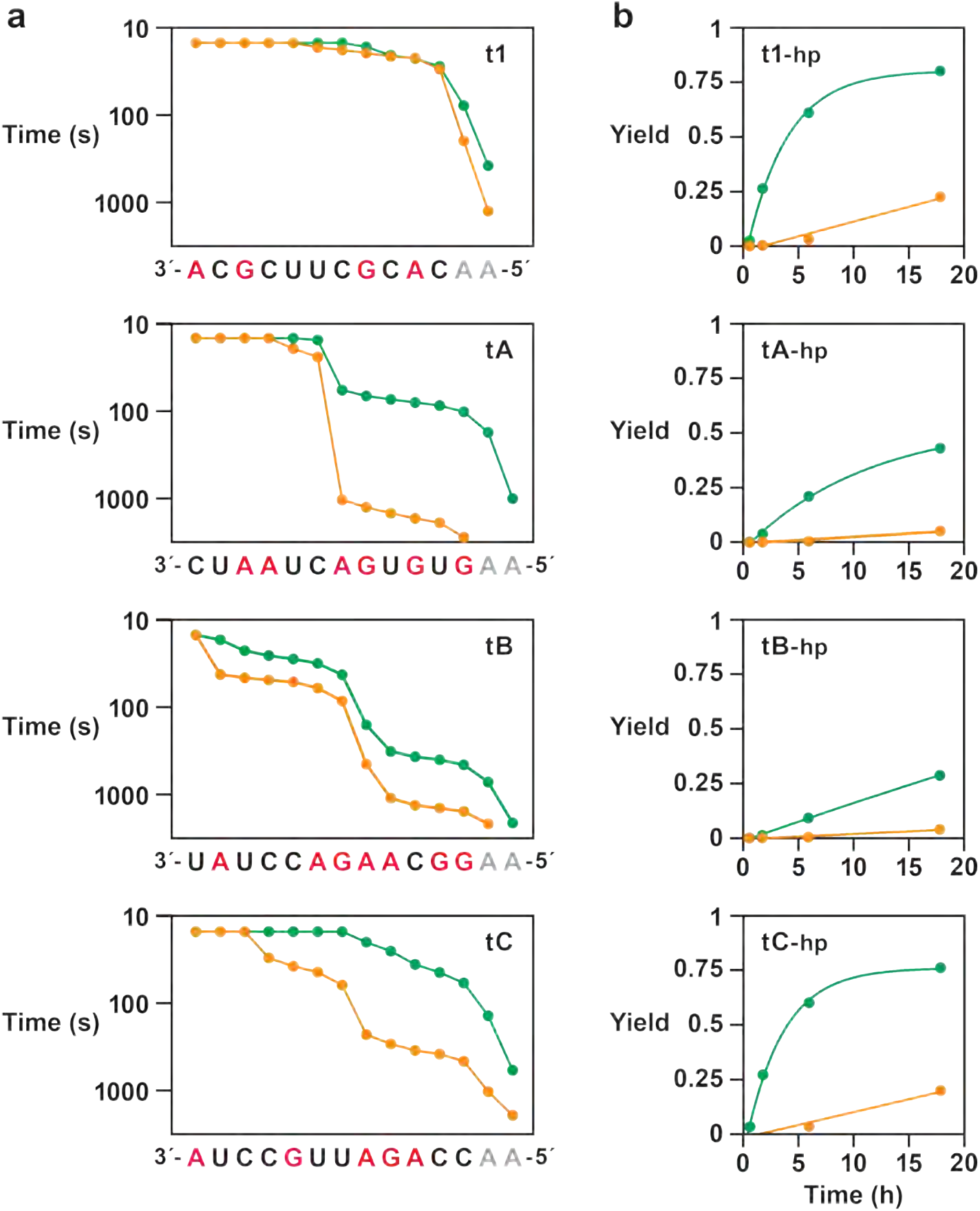
Improved sequence generality. (a) Activity of the 52-2 (orange) and 85h34 (green) polymerases on four short RNA templates of varying sequence (purine nucleotides in red; spacer nucleotides in gray). Time (log scale) is the number of seconds needed to achieve ≥15% yield at the indicated nucleotide position or longer. (b) Activity on four corresponding templates with an added 6-base-pair hairpin structure at the 5’ end. Yield refers to full-length products, including copying the hairpin. Reaction conditions: 100 nM polymerase, 100 nM template, 80 nM primer, 50 mM MgCl_2_, and 2 mM NTPs at pH 8.3 and 25 °C.

Tandem repeats of simple unstructured template sequences have been used to gauge the ability of the polymerase to produce long RNAs, most notably tandem repeats of **t1**. For example, the tC19Z polymerase is able to copy 5–10 tandem repeats of **t1** in 24 h,^6^ while 52-2 can copy 5 repeats in 10 min.^14^ 52-2 and 85h34 perform similarly on this 5-fold repeat template in the presence of 50 mM Mg^2+^, yielding full-length products in 10 min (Table S3). However, on tandem repeats of the more heterogeneous sequences, 52-2 requires 100 min to complete the synthesis of (tC)_5_ and is unable to complete the other two syntheses. In contrast, 85h34 completes all three syntheses in 10–100 min.

The ability to copy through structured regions of the template has also improved. The capacity to unwind and copy stem structures within the template was first observed for the 24-3 polymerase,^7^ and subsequent evolved polymerases have improved upon this ability as selection required the synthesis of more complex and structured ribozymes. For each of the four templates, **t1**, **tA**, **tB**, and **tC**, a six base-pair hairpin closed by a UACG tetraloop was installed at the 5’ end and the 52-2 and 85h34 polymerases were compared for their ability to synthesize full-length products in the presence of 50 mM Mg^2+^. 85h34 can copy through each hairpin in 2 h and reaches 20–80% yield of full-length products after 20 h, whereas 52-2 requires 18 h to copy through the hairpins and does so in much lower yield (Figure 4b).

### Synthesis of Functional RNAs

In view of the improved yield of RNA-catalyzed synthesis of the class I ligase and evidence for improved generality with regard to template sequence and structure, the 85h34 polymerase was challenged to synthesize various functional RNAs in comparison to the 52-2 polymerase. Although 52-2 performs optimally at 200 mM Mg^2+^, its fidelity is compromised under this condition,^14,15^ as is that of the population of polymerases at round 85. Thus the 85h34 and 52-2 polymerases were assessed at 50 mM Mg^2+^ and either 17 °C for 52-2 or 25 °C for 85h34. The lower temperature for 52-2 was found to be necessary to obtain a detectable yield of long primer-extension products such as the class I ligase after 24 h, whereas no full-length products are obtained at 25 °C.

The two polymerases were challenged to synthesize four different functional RNAs: the Mango aptamer (28 nucleotides), hammerhead ribozyme (33 nucleotides), F1 ligase (49 nucleotides), and class I ligase (97 nucleotides) (Figure 5). These RNAs span a range of sizes, structural complexity, and sequence composition, providing a good test of the generality of polymerase activity. The 85h34 polymerase synthesizes the Mango aptamer in 40% yield after 24 h, compared to <10% yield for 52-2. Both polymerases achieve ∼25% yield for the hammerhead ribozyme and ∼10% yield for the F1 ligase after 24 h. The difference is more dramatic for the class I ligase, which 85h34 produces in 12.5% yield after 24 h, compared to <1% yield for 52-2. The Mango aptamer and class I ligase are both challenging targets — the aptamer because it has few guanosine residues at its 3’ end and therefore few cytosine residues in the corresponding portion of the template, and the class I ligase because the template has extensive secondary structure that must be displaced by the polymerase in order to yield a full-length copy.

**Figure 5.**
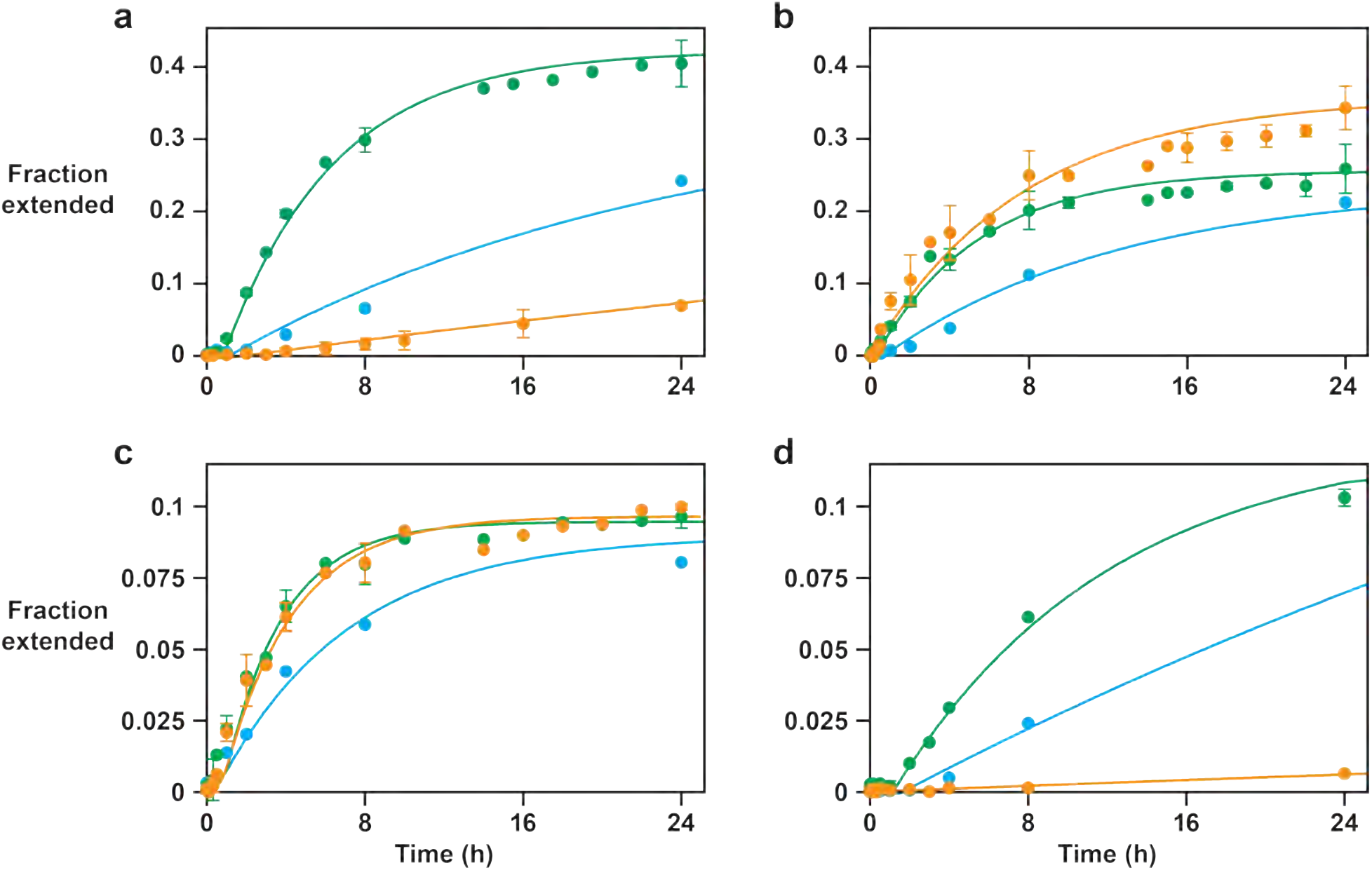
RNA-catalyzed synthesis of functional RNAs. Time course for synthesis of (a) Mango aptamer, (b) hammerhead, (c) F1 ligase, and (d) class I ligase, catalyzed by either the 52-2 polymerase in the presence of: i) 50 mM MgCl_2_ and 2 mM NTPs at 17 °C (orange); or the 85h34 polymerase in the presence of either: ii) 50 mM MgCl_2_ and 2 mM NTPs at 25 °C (green); or iii) 10 mM MgCl_2_, 0.5 mM NTPs, and 8% PEG at 27 °C (blue), all at pH 8.3. Reactions were performed in at least triplicate, with error bars representing standard error.

As noted above, the 85h34 polymerase is able to synthesize the class I ligase in the presence of only 10 mM Mg^2+^, together with 8% (w/v) PEG8000 as a molecular crowding reagent. This is the lowest concentration of Mg^2+^ cofactor ever used by a polymerase ribozyme to copy RNA. The class I ligase and all prior forms of the polymerase through round 52 were evolved to operate in the presence of 200 mM MgCl_2_. The ability of 85h34 to synthesize each of the four functional RNAs was evaluated at 10 mM Mg^2+^, which resulted in yields after 24 h that were >75% of what is obtained at 50 mM Mg^2+^.

Although the production of full-length RNAs by the polymerase entails many sequential single-nucleotide additions, the observed rate of synthesis by both 52-2 and 85h34 fits well to a single-exponential rise to maximum, consistent with first-order kinetics. This observation is likely due to the dominance of template regions that are the most difficult to copy in defining the rate-limiting step for synthesis of full-length products. As such, it is notable that the ribozymes that are synthesized most efficiently by the 52-2 polymerase at 50 mM Mg^2+^, the hammerhead and F1 ligase, are produced with only a slightly faster rate by 85h34. For synthesis of the hammerhead ribozyme, the observed reaction rates are 0.13 and 0.18 h^−1^ for 52-2 and 85h34, respectively; for synthesis of the F1 ligase, the rates are 0.28 and 0.33 h^−1^, respectively. These results are consistent with the earlier finding that both ribozymes perform well on unstructured, C-rich templates, but that 85h34 excels on more difficult templates. For the more difficult syntheses of the Mango aptamer and class I ligase, 85h34 outperforms 52-2 by 20- and 13-fold, respectively.

### Analysis of Polymerase Fidelity

Preliminary analysis revealed that the 85h34 polymerase synthesizes the full-length class I ligase with significantly fewer mutations compared to 52-2, as would be expected given the selection pressures that were applied during evolution. To investigate the distribution of mutations generated by the two polymerases, the products of RNA-catalyzed synthesis of the hammerhead, F1 ligase, and class I ligase were analyzed by deep sequencing for reactions carried out at 50 mM Mg^2+^ for both polymerases and at 10 mM Mg^2+^ for 85h34. The reaction times were chosen so that the extent of reaction was comparable in all cases, corresponding to ∼1% yield of full-length products relative to the starting amount of primer. The fidelity per nucleotide of the 52-2 polymerase decreases with increasing product length, achieving 93% fidelity for the hammerhead ribozyme, but decreasing to 91% for the F1 ligase and 86% for the class I ligase (Figure 6a). In contrast, the fidelity of 85h34 is consistent over the three different product lengths, with fidelities of 91%, 93%, and 91%, respectively, for synthesis of the three ribozymes at 50 mM Mg^2+^, and 94%, 95%, and 94%, respectively, at 10 mM Mg^2+^.

**Figure 6.**
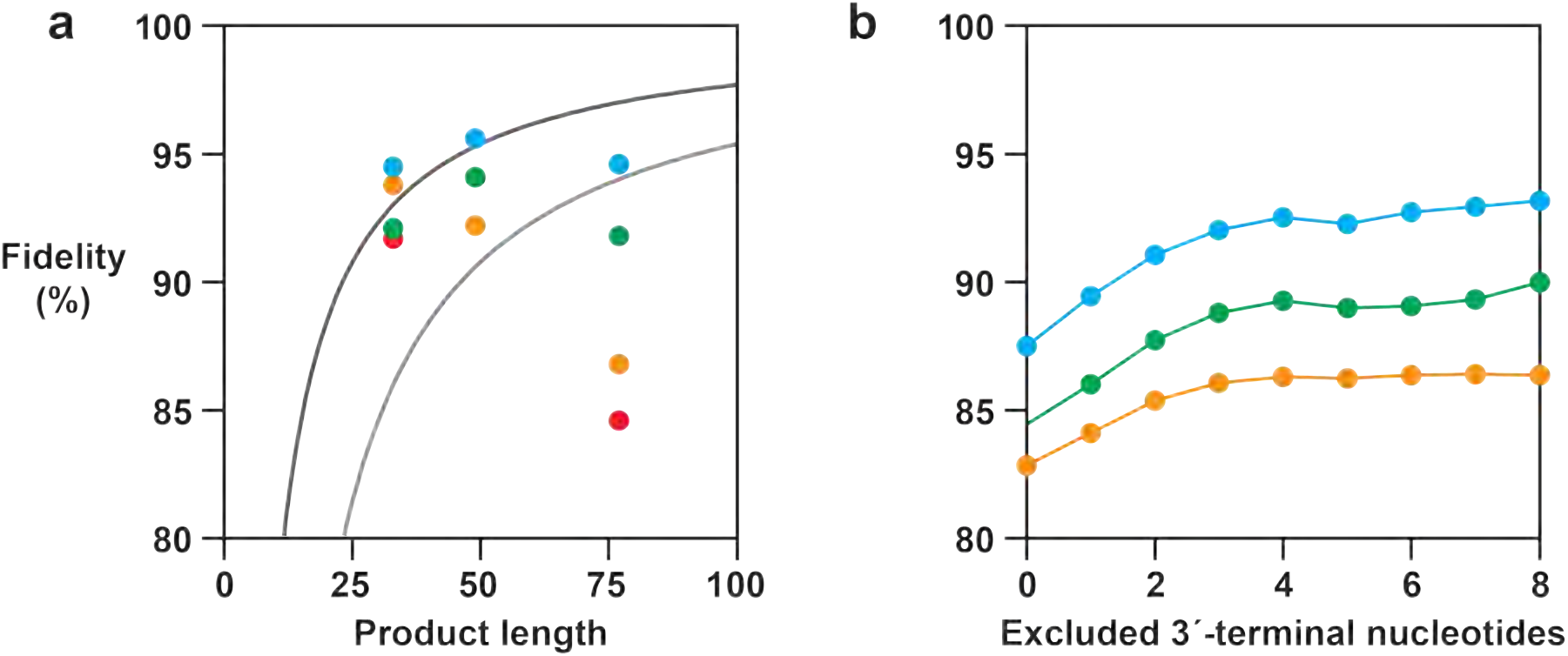
Fidelity of RNA synthesis. (a) Per-nucleotide fidelity of the 52-2 polymerase in the presence of: i) 200 mM MgCl_2_ and 4 mM NTPs at 17 °C (red); or ii) 50 mM MgCl_2_ and 2 mM NTPs at 17 °C (orange); and of the 85h34 polymerase in the presence of: iii) 50 mM MgCl_2_ and 2 mM NTPs at 25 °C (green); or iv) 10 mM MgCl_2_, 0.5 mM NTPs, and 8% PEG at 27 °C (blue); all at pH 8.3. The synthesized products are the hammerhead (33 nucleotides), F1 ligase (49 nucleotides), and class I ligase (77 nucleotides). Curved lines indicate the error threshold as a function of product length,^9^ assuming 25% neutral positions and a fitness advantage for the most advantageous species of either 10-or 100-fold (dark and light gray lines, respectively). (b) Position-specific fidelity of synthesis, including both partial- and full-length products, as a function of the distance of the nucleotide from the 3’ end of the product. Fidelity values refer to all nucleotides that are further from the 3’ terminus than the specified number of excluded nucleotides.

It was previously observed that polymerase fidelity decreases with longer reaction times and that the frequency of mutations is higher near the 3’ end of the extension products.^13^ These observations suggest that the polymerase is prone to stalling following incorporation of mutations, a phenomenon that has also observed for polymerase proteins.^25,26^ Stalling delays further extension, but with long reaction times these intermediates may eventually be extended to yield full-length products that contain the stall-inducing mutations. More challenging templates that require a longer time to copy provide more opportunities for polymerases that are stalled at a site of mutation to resume synthesis, thereby decreasing overall fidelity. To explore how these effects contribute to the fidelity of the 52-2 and 85h34 polymerases, all partial- and full-length extension products were included in the analysis. The depth of sequencing was 40 million reads per sample to ensure that even rare extension products would be sampled, providing fidelity measurements that are accurate to 1%.

This analysis revealed that for both 52-2 and 85h34 fidelity is lower when considering both partial- and full-length products, but rises as positions near the 3’ terminus are excluded from the analysis (Figure 6b). The average fidelity across all nucleotide positions reflects the insertional fidelity of the polymerase, the probability that an incorrect nucleotide is incorporated regardless of subsequent effects of polymerase stalling. The fidelity plateau that is reached when excluding the first four positions upstream of the 3’ terminus represents the effective fidelity after allowing for polymerase stalling at mismatched termini. The difference in fidelity between the 52-2 and 85h34 polymerases at 50 mM Mg^2+^ is largely due to enhanced discrimination by 85h34 against extending terminal mismatches. The modest difference in insertional fidelities of 82.8% and 84.5% for 52-2 and 85h34, respectively, becomes more substantial when considering only internal positions, in that case 86.3% and 90.8%, respectively. In contrast, the further gain in fidelity when 85h34 operates at 10 rather than 50 mM Mg^2+^ appears to be driven by both improved insertional fidelity, 87.5% at 10 vs. 84.5% at 50 mM Mg^2+^, and a similar improvement when considering only internal positions, 94.0% at 10 vs. 90.8% at 50 mM Mg^2+^. At both Mg^2+^concentrations the increase in fidelity for 85h34 due to decreased extension of mismatched termini is 5%.

The most common mutational errors made by the two polymerases are G:U wobble mismatches, misincorporation opposite template adenosine positions, and single-nucleotide insertions and deletions. The frequency of all three types of mutations is lower for 85h34 compared to 52-2 when measured in the presence of 50 mM Mg^2+^, and lower still for 85h34 at 10 mM Mg^2+^ (Table S4). There are idiosyncratic effects on fidelity for specific nucleotide positions, but 85h34 consistently outperforms 52-2, especially in situations where 52-2 performs poorly. In general, nucleotide positions that are the most difficult to copy accurately are those where 85h34 shows the greatest improvement in fidelity.

## Discussion

The ability of RNA to catalyze the RNA-templated polymerization of NTPs has been established for 30 years.^27^ Efforts to use directed evolution to enhance this activity began a few years later,^11^ and have continued in multiple laboratories, leading to polymerase ribozymes that can synthesize substantially longer RNA products.^6,28^ Product length is an important measure of polymerase activity, but with the aim of propagating heritable information through the RNA-catalyzed evolution of RNA, sequence generality and copying fidelity are also critical factors.

Broad sequence generality is important for access to “sequence space”, the set of possible sequences that the evolving population can explore. If there are substantial restrictions on the nucleotide composition of templates that can be copied, then the reciprocal synthesis of complementary strands will be impaired. For example, the need for C-rich template sequences will result in G-rich product strands that are difficult to copy. Ideally, all possible sequences can be copied with equal efficiency, but even the most sophisticated protein polymerases have some bias with regard to template sequence.^29,30^ Sequence generality also pertains to copying structured template regions, which are inevitable for RNAs of heterogenous sequence, especially for templates that encode stem elements that are critical to the structure of functional RNAs. The complement of a stem is also a stem, so such elements need to be tolerated within both the template and product strands.

Copying fidelity is fundamental to information transfer across multiple cycles of reciprocal strand synthesis. Occasional copying errors introduce variation into the evolving population, which are culled by selection to provide the opportunity for evolutionary innovation. But if the error frequency is above a critical threshold, there is a cascade of errors that cannot be contained by selection, and the population degenerates into non-functional sequences.^8^ The prior 71-89 polymerase ribozyme was the first ribozyme with sufficient fidelity to be able to replicate and evolve another ribozyme, the 33-nucleotide hammerhead.^16^ The higher the fidelity, the larger the amount of functional information that can be evolved. Ultimately, the notion of the RNA world turns on the possibility of an RNA polymerase ribozyme with sufficient sequence generality and copying fidelity that it can evolve copies of itself.

Over the course of 52 prior rounds of directed evolution, the polymerase ribozyme became more adept with regard to catalytic efficiency and sequence generality, enabling it to synthesize ever larger functional RNAs. However, these improvements came with a trade-off of reduced accuracy due to the low information content of the functional RNAs that the polymerase was required to synthesize. Reduced fidelity is most apparent when the polymerase is pushed to its limits by synthesizing the large and complex class I ligase, resulting in product RNAs with a high level of mutations that render most molecules inactive. Nonetheless, the ability to synthesize the class I ligase made it possible to carry out further rounds of evolution in which the polymerase was required to synthesize this more complex functional RNA, thus placing strong selection pressure to favor improvements in sequence generality and copying fidelity. The improved catalytic efficiency achieved by round 52 also made it possible to reduce the concentration of Mg^2+^, further favoring improved fidelity. The 85h34 polymerase has substantially improved sequence generality and copying fidelity compared to the 52-2 polymerase, and now provides the opportunity to impose even more stringent selection constraints. This bootstrapping process may ultimately provide a path to the realization of a self-evolving RNA.

The population dynamics over the course of evolution were investigated through deep sequencing, cluster analysis, and phylogenetic reconstruction, revealing the emergence and competition of various clades with distinct combinations of mutations. Following an initial burst of new haplotypes at the outset of selection for synthesis of the class I ligase, four persistent clades arose over the subsequent 33 rounds. An early dominant clade, 58b, exhibited high fidelity but reduced activity, and was unable to persist when the polymerases were required to complete the synthesis of the class I ligase in shorter times and at lower Mg^2+^ concentrations. Two clades that had been present at low frequency, 58a and 52b, then rose to displace 58b and became further differentiated from the 52-2 polymerase by accumulating additional mutations. The P6 and P8 stems were remodeled in clade 58a and the P4 and A4 stems were remodeled in clade 52b, in each case by inverting Watson-Crick pairs. However, because the two strands that comprise the P6, P8, and P4 stems are well separated in the primary sequence of the ribozyme, productive recombination between the two clades was unlikely. Experimenter intervention came to the rescue, combining advantageous traits from 58a and 52b in a hybrid molecule, the 85h34 polymerase, to provide the most general and accurate RNA polymerase ribozyme to date.

Deep sequencing was used to examine the fidelity of the 85h34 polymerase in more detail. The overall fidelity is in the range of 91–93% per nucleotide for reactions carried out at 50 mM Mg^2+^ and 94–95% per nucleotide at 10 mM Mg^2+^. This is a substantial improvement compared to the behavior of the 52-2 polymerase, and is especially the case for templates that are longer and more difficult to copy. The improved fidelity of 85h34 is due both to reduced incorporation of mismatched nucleotides and a reduced propensity to extend mismatched termini. The lower amount of misincorporation is more pronounced at the lower concentration of Mg^2+^, whereas the lower propensity to extend mismatch termini is seen at both Mg^2+^ concentrations.

In a parallel study to this one, the 85h34 polymerase proved amenable to structural analysis by cryo-electron microscopy. A 2.8 Å resolution structure was obtained, revealing how adaptions over the course of evolution resulted in a well-defined active site that positions the template-bound primer and incoming NTP for catalysis.^24^ Going forward, such knowledge of the polymerase structure, together with the powerful search process of directed evolution, provides an opportunity for further improvement of the fidelity and sequence generality of the polymerase ribozyme.

## ASSOCIATED CONTENT

### Supporting Information

The Supplementary Information is available free of charge at: <URL>

## AUTHOR INFORMATION

### Author Contributions

D.P.H., and G.F.J. designed research; D.P.H. and S.N. performed research; D.P.H., S.N., and G.F.J. analyzed data; and D.P.H. and G.F.J. wrote the paper.

### Funding

This work was supported by NASA grant 80NSSC22K0973 and Sloan Foundation grant 25294.

### Notes

The authors declare no competing financial interest.

## ACKNOWLEDGMENT

The authors are grateful to Dmitry Lyumkis for helpful comments on the manuscript and to the members of the Simons Collaboration on the Origins Life, supported by the Simons Foundation, for many stimulating discussions over the years.

## Supporting Information

**Table S1.**
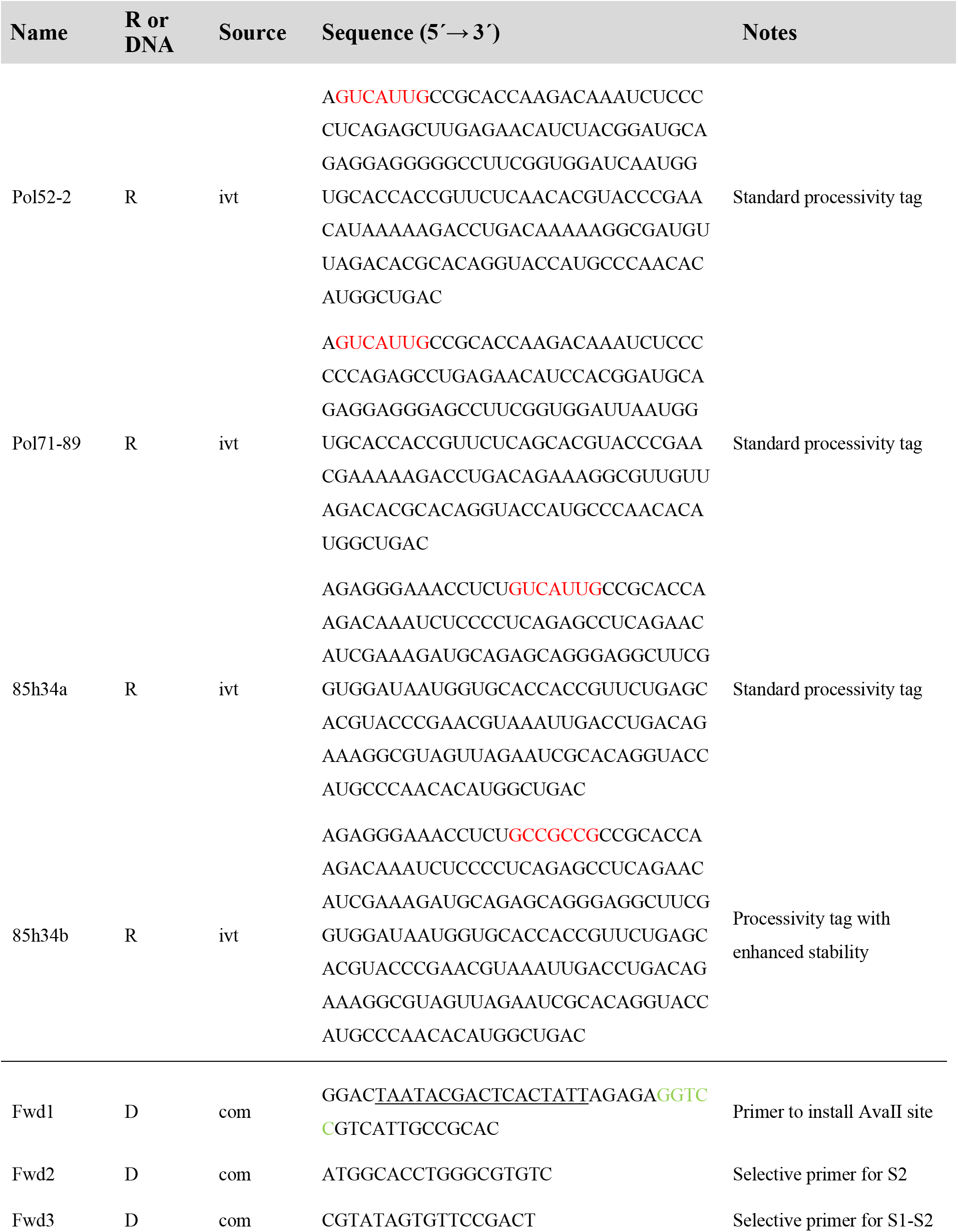
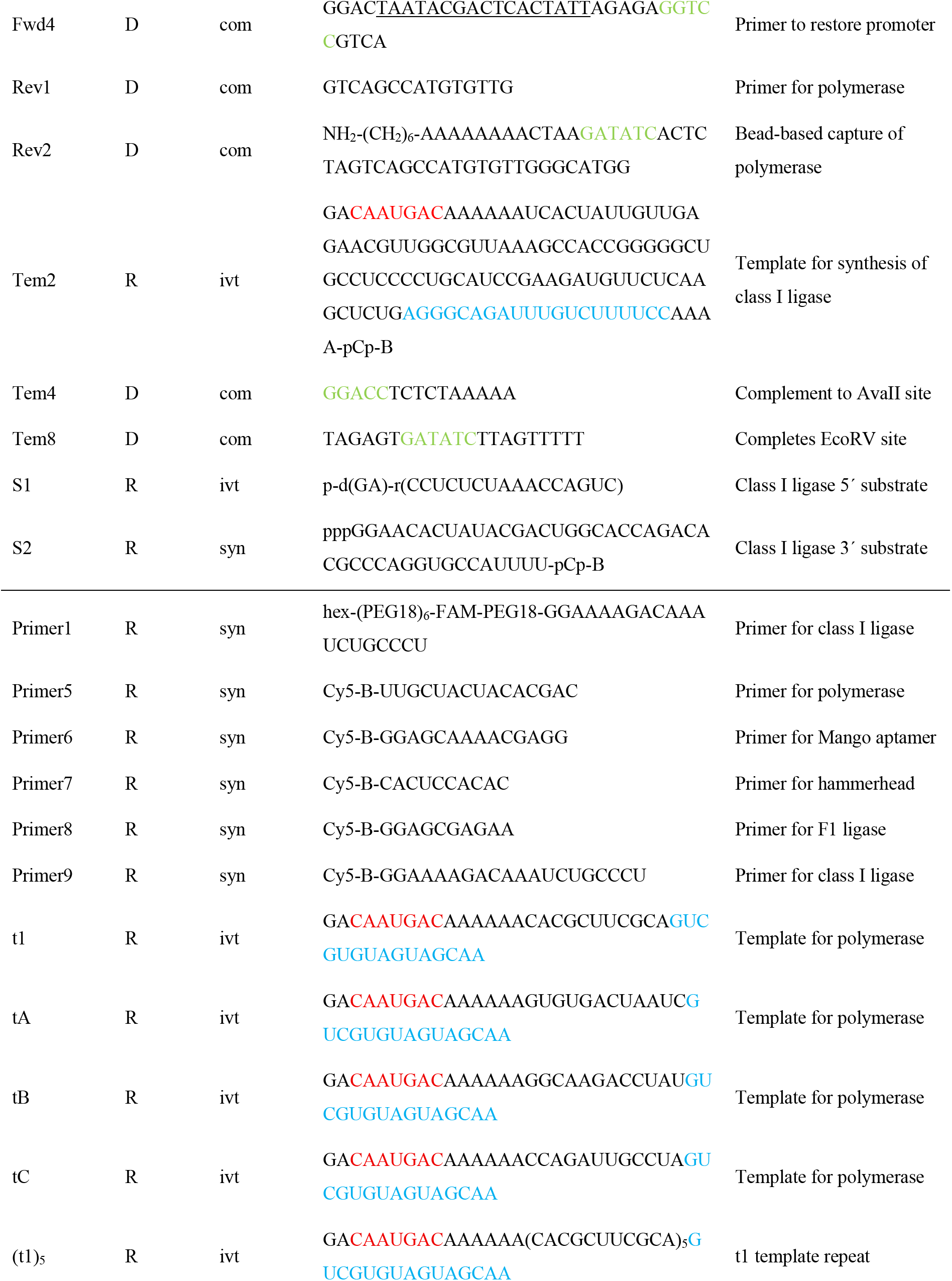
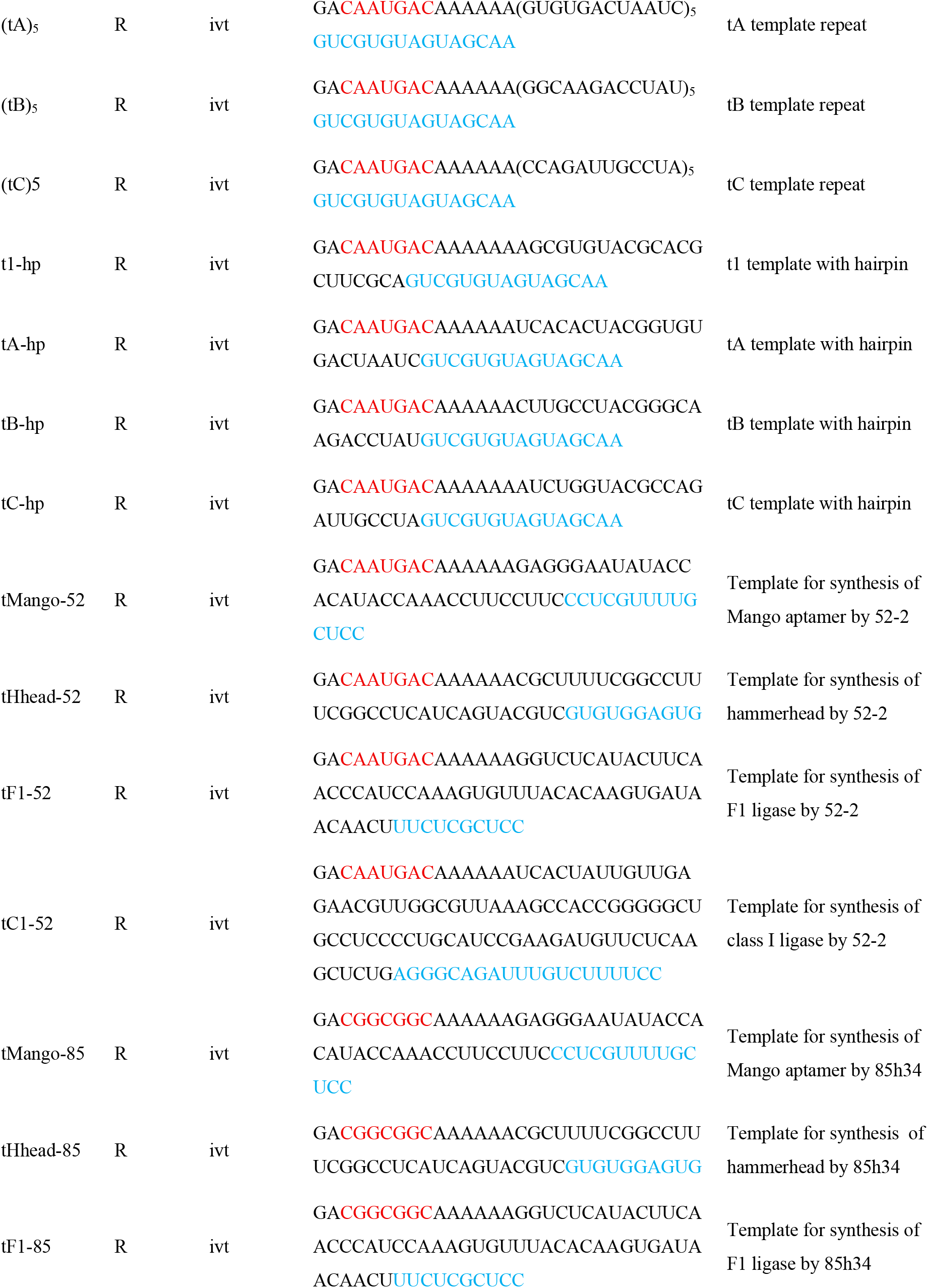
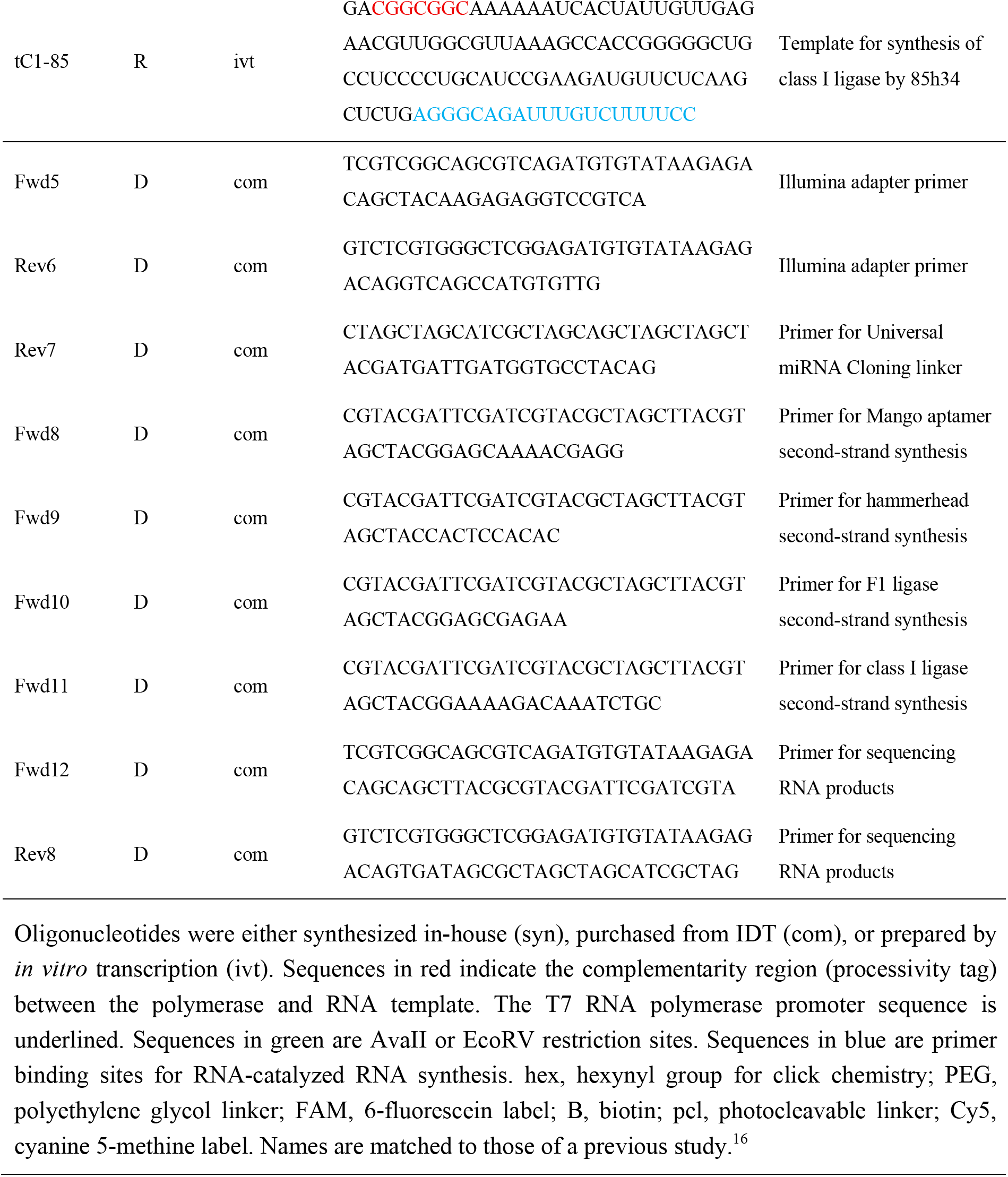
Sequences of RNA and DNA molecules used in this study.

**Table S2.** Conditions During Each Round of Evolution.

| Round | Time (h) | [MgCl<sub>2</sub>] (mM) | Mutagenesis | Sequencing |
| --- | --- | --- | --- | --- |
| 53 | 24 | 200 | + |  |
| 54 | 24 | 200 | + |  |
| 55 | 24 | 200 | + |  |
| 56 | 24 | 200 | + |  |
| 57 | 24 | 200 | + |  |
| 58 | 24 | 200 | + | + |
| 59 | 24 | 200 |  |  |
| 60 | 24 | 200 | + | + |
| 61 | 24 | 200 | + | + |
| 62 | 5 | 100 |  | + |
| 63 | 2.5 | 100 | + |  |
| 64 | 1 | 100 |  |  |
| 65 | 1 | 100 |  |  |
| 66 | 1 | 100 |  |  |
| 67 | 1 | 100 | + | + |
| 68 | 1 | 100 | + |  |
| 69 | 1 | 100 | + |  |
| 70 | 1 | 100 | + |  |
| 71 | 1 | 83 | + |  |
| 72 | 1 | 69 |  | + |
| 73 | 1 | 69 | + |  |
| 74 | 1 | 66 |  |  |
| 75 | 1 | 66 |  | + |
| 76 | 1 | 66 |  |  |
| 77 | 1 | 66 |  |  |
| 78 | 1 | 66 | + |  |
| 79 | 1 | 66 |  | + |
| 80 | 1 | 66 |  |  |
| 81 | 1 | 66 |  |  |
| 82 | 1 | 66 |  |  |
| 83 | 1 | 66 | + | + |
| 84 | 1 | 66 |  | + |
| 85 | 1 | 50 |  | + |

Reaction time and concentration of MgCl<sub>2</sub> for RNA-catalyzed synthesis of the class I ligase. Mutagenic PCR was carried out during completion of the indicated rounds. Deep sequencing of the population was carried out as indicated, prior to any PCR mutagenesis.

**Table S3.**
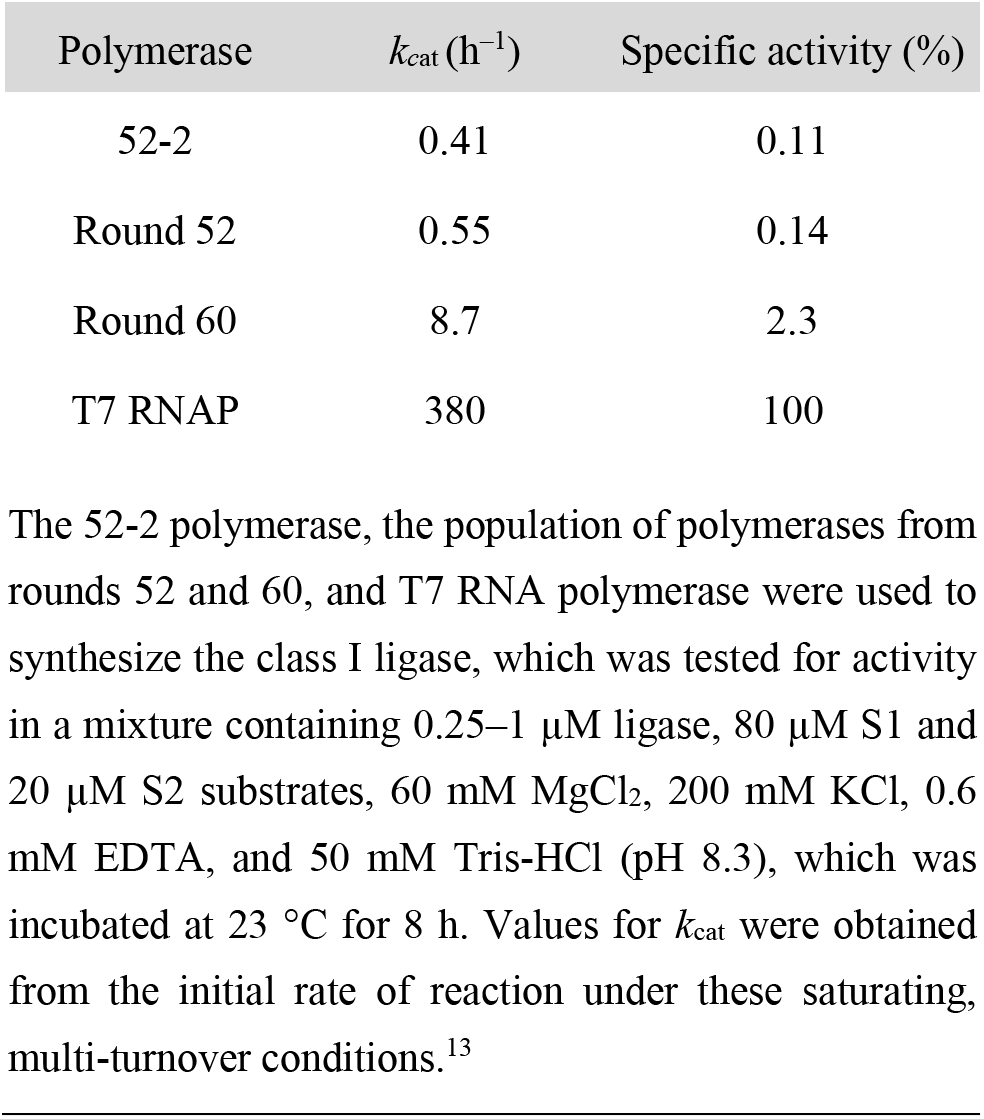
Activity of Class I Ligase Products.

| Polymerase | $k_{\text{cat}}$ (h<sup>-1</sup>) | Specific activity (%) |
| --- | --- | --- |
| 52-2 | 0.41 | 0.11 |
| Round 52 | 0.55 | 0.14 |
| Round 60 | 8.7 | 2.3 |
| T7 RNAP | 380 | 100 |

The 52-2 polymerase, the population of polymerases from rounds 52 and 60, and T7 RNA polymerase were used to synthesize the class I ligase, which was tested for activity in a mixture containing 0.25–1 $\mu\text{M}$ ligase, 80 $\mu\text{M}$ S1 and 20 $\mu\text{M}$ S2 substrates, 60 mM $\text{MgCl}_2$ , 200 mM KCl, 0.6 mM EDTA, and 50 mM Tris-HCl (pH 8.3), which was incubated at 23 °C for 8 h. Values for $k_{\text{cat}}$ were obtained from the initial rate of reaction under these saturating, multi-turnover conditions.<sup>13</sup>

**Table S4.** Contribution to Infidelity by Major Classes of Mutations.

| Polymerase | [MgCl<sub>2</sub>] (mM) | Fidelity (%) | Wobble (%) | U→V (%) | Indel (%) |
| --- | --- | --- | --- | --- | --- |
| 52-2 | 50 | 86.3 | 4.0 | 3.3 | 5.4 |
| 85h34 | 50 | 89.3 | 3.1 | 2.7 | 4.7 |
| 85h34 | 10 | 92.5 | 2.4 | 1.6 | 3.4 |

Fidelity of all partial- and full-length products for positions at least four nucleotides from the 3' terminus for RNA-catalyzed synthesis of the hammerhead, F1 ligase, and class I ligase. Wobble mutations are A→G or C→U substitutions, U→V mutations are substitution of U by any other nucleotide, and indels are both insertions and deletions. Fidelity was calculated as the geometric average across all four expected nucleotides. Insertion rates were calculated separately and added to the contribution to fidelity from deletions.

**Figure S1.**
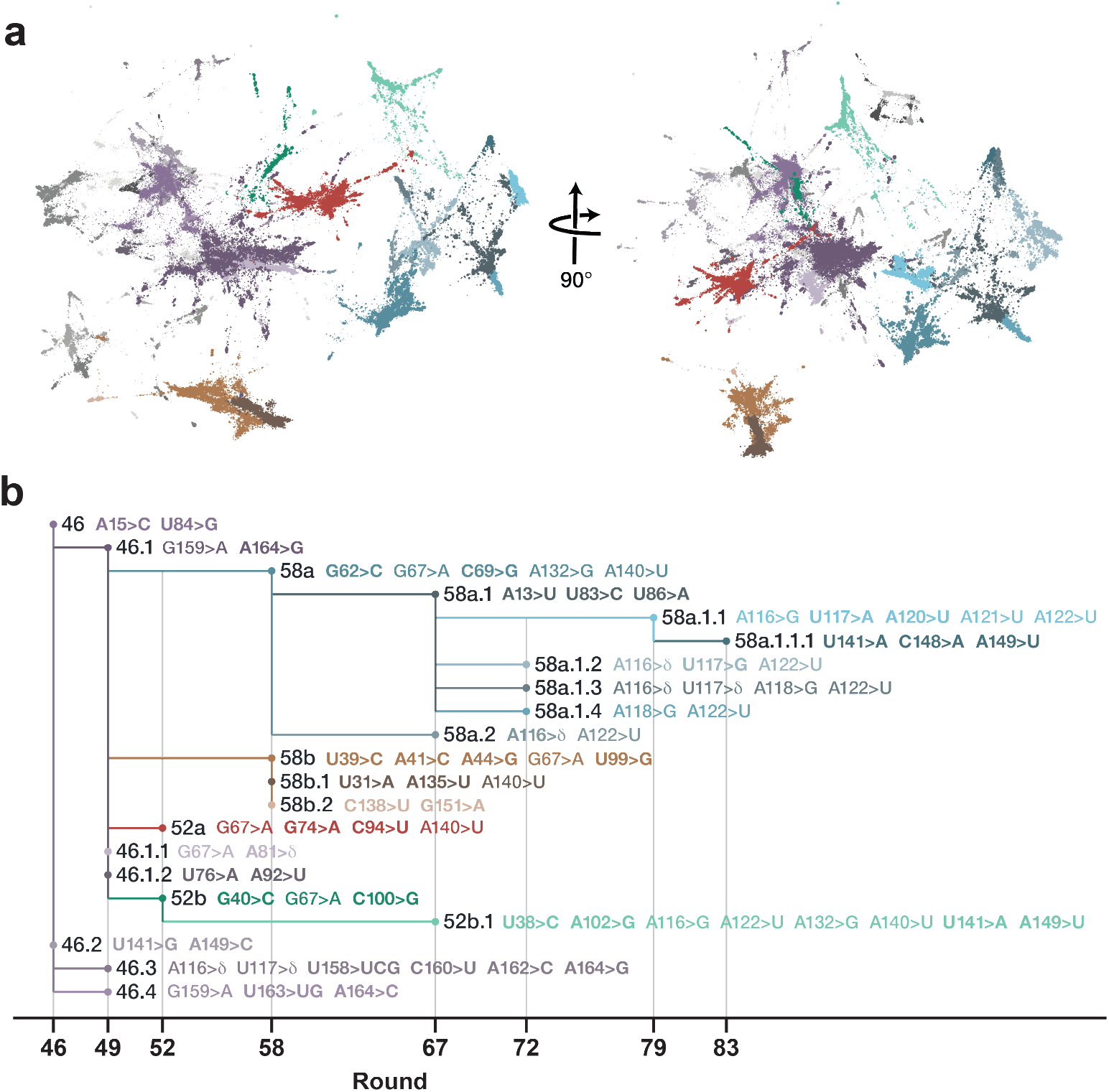
Population dynamics over the course of evolution. (a) Three-dimensional embedding of the distribution of polymerase sequences in rounds 39–85, depicting the most common clusters, linked by filaments of transitional forms. Colors correspond to those shown in Figure 2b. Clades that were present prior to the emergence of clade 46 are shown in gray. (b) Phylogenetic tree built from the clusters by maximum parsimony. Individual clades that reached >20% of the population in at least one round are named based on the round in which they first appeared. Differentiating mutations for each clade are listed, with those that are unique to the clade shown in bold.

**Figure S2.**
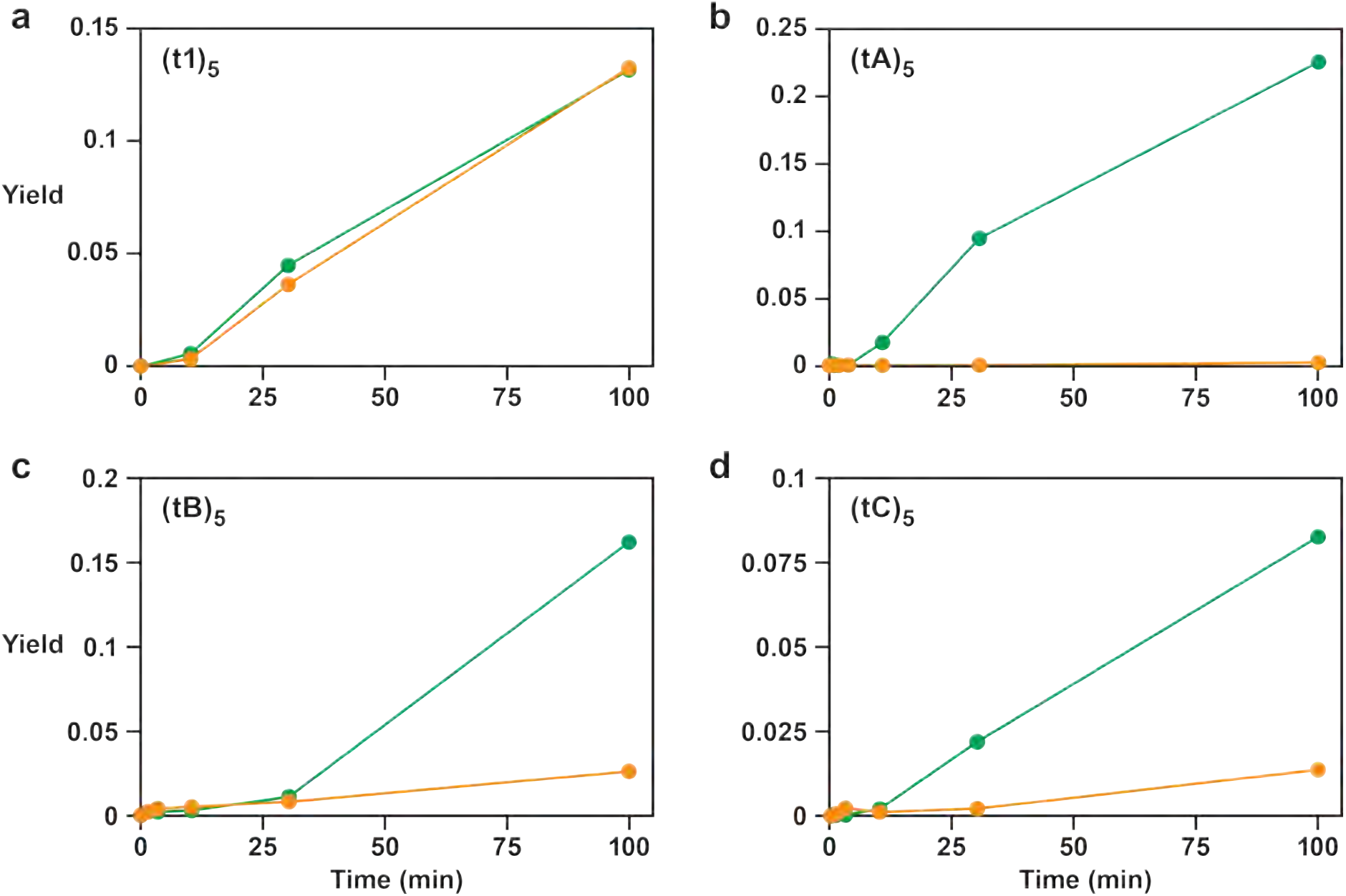
RNA-catalyzed copying of repeat templates. (a) Activity of the 52-2 (orange) and 85h34 (green) polymerases on five tandem repeats of each of the four short RNA templates of varying sequence (t1, tA, tB, and tC). Yield refers to completion of the fifth repeat along the template. Reaction conditions: 100 nM polymerase, 100 nM template, 80 nM primer, 50 mM MgCl_2_, and 2 mM NTPs at pH 8.3 and 25 °C.

## Notes

### Competing Interest Statement

The authors have declared no competing interest.

